# Single cell RNA-seq study of wild type and Hox9,10,11 mutant developing uterus

**DOI:** 10.1101/395574

**Authors:** Michael L. Mucenski, Robert Mahoney, Mike Adam, Andrew S. Potter, S. Steven Potter

## Abstract

The uterus is a remarkable organ that must guard against infections while maintaining the ability to support growth of a fetus without rejection. The *Hoxa10* and *Hoxa11* genes have previously been shown to play essential roles in uterus development and function. In this report we show that the *Hoxc9,10,11* genes play a redundant role in the formation of uterine glands. In addition, we use single cell RNA-seq to create a high resolution gene expression atlas of the developing wild type mouse uterus. Cell types and subtypes are defined, for example dividing endothelial cells into arterial, venous, capillary, and lymphatic, while epithelial cells separate into luminal and glandular subtypes. Further, a surprising heterogeneity of stromal and myocyte cell types are identified. Transcription factor codes and ligand/receptor interactions are characterized. We also used single cell RNA-seq to globally define the altered gene expression patterns in all developing uterus cell types for two Hox mutants, with 8 or 9 mutant Hox genes. The mutants show a striking disruption of Wnt signaling as well as the Cxcl12/Cxcr4 ligand/receptor axis.

**Summary statement:** A single cell RNA-seq study of the developing mouse uterus defines cellular heterogeneities, lineage specific gene expression programs and perturbed pathways in Hox9,10,11 mutants.

## Introduction

The uterus must guard against infections while receiving an allograft implant, the embryo, without rejection. It is a dynamic tissue with cyclic developmental changes, as well as responses to steroids that lead to receptivity for implantation. Proper uterus function is required for fertility, and disorders can lead to endometriosis and neoplasia. At birth, the uterus is composed of simple epithelium surrounded by undifferentiated mesenchyme. The uterus then differentiates into a columnar luminal epithelium (LE), surrounded by stroma, which in turn is surrounded by two myometrial layers (Hayashi et al., 2011). Uterine glands secrete LIF and calcitonin, each required for fertility (Stewart et al., 1992; Zhu et al., 1998). Uterine gland formation in the mouse begins by post-natal day (PND) 6 with the invagination or budding of the LE to form glandular epithelium (GE)(Bigsby et al., 1990; Cooke et al., 2012). By PND12 uterine endometrial glands extend from the LE into the surrounding stroma and the longitudinal layer of the myometrium is organized into bundles of smooth muscle cells (Cunha et al., 1992). Gland development is a continuous process that extends beyond puberty (Brody and Cunha, 1989; Stewart et al., 2011).

Hox genes are known to play important roles in uterus development and function. There are thirty nine mammalian Hox genes, arranged in four clusters located on four separate chromosomes. The Hox genes of these HoxA, B, C, and D clusters are classified into 13 paralogous groups based on sequence similarity. The study of Hox genes is confounded by their extensive functional overlap. While the paralogous Hox genes show the greatest functional similarity, there is also extensive evidence for shared functions of Hox genes that lie near each other on a cluster (Branford et al., 2000; Davis et al., 1995; Favier et al., 1996; Greer et al., 2000; Zhao and Potter, 2001). Of interest, the 16 most 5’ Hox genes of paralog groups 9-13 are quite closely related and are designated Abd-B type Hox genes. The Hox9,10,11 paralog genes within this group are particularly closely related. Early studies showed that the *Hoxa10* and *Hoxa11* genes play key roles in the development and function of the female reproductive tract. Homozygous mutation of either of these Hox genes results in partial homeotic transformation of the uterus to the more anterior oviduct and significantly reduced fertility due to perturbed uterus function (Benson et al., 1996; Gendron et al., 1997; Hsieh-Li et al., 1995; Lim et al., 1999; Ma et al., 1998; Satokata et al., 1995). *Hoxa10* mutation results in defective implantation and decidualization, resulting in reduced fertility (Das, 2010). *Hoxa10* is expressed in the luminal and glandular epithelium on days 1 and 2 of pregnancy, expands to stroma on day 3 and is restricted to stroma on day 4 (Das, 2010). Mutants show reduced stromal proliferation in response to estrogen and progesterone.

Of interest, while the *Hoxa10* and *Hoxa11* genes have defined functions in female fertility, single homozygous mutation of the paralogous *Hoxc10, Hoxd10, Hoxc11* and *Hoxd11* genes gave no reported infertility. Further, the closely related Hox9 paralog genes could be mutated in combination, such as *Hoxa9-/- Hoxb9-/- Hoxd9-/-*, with the resulting triple homozygous mutant mice having normal uterus function, although their mammary glands showed defective development post pregnancy (Chen and Capecchi, 1999). Mice with multiple combinations of mutations of either the Hox10 or Hox11 paralog groups have interesting kidney and limb abnormalities, but no described infertility phenotypes extending beyond those associated with the *Hoxa10* and *Hoxa11* genes (Wellik and Capecchi, 2003; Wellik et al., 2002). These results suggest unique roles for *Hoxa10* and *Hoxa11* in uterus development and function.

We have, however, previously shown that it is possible to identify uterine functions for other paralogous Hox9,10,11 genes through the use of a “sensitized” genotype that includes reduced *Hoxa10* and *Hoxa11* activity. For example, female *Hoxd9,10,11-/-* mice (with simultaneous homozygous mutation of *Hoxd9, Hoxd10* and *Hoxd11*) give normal size litters, as do *Hoxa9,10,11+/-* mice, while mice that are both *Hoxa9,10,11+/-* and *Hoxd9,10,11-/-* are completely infertile (Raines et al., 2013). The *Hoxd9,10,11* genes have redundant function with *Hoxa9,10,11* in oviduct/uterus identity determination and also have key roles in uterine immune and noncoding RNA gene regulation (Raines et al., 2013).

In this report we extend this approach to search for possible female fertility functions for the *Hoxc9,10,11* genes. We observed that while *Hoxa9,10,11*+/-, *Hoxd9,10,11*+/- mice showed reduced fertility, with about half normal litter size, mice that were in addition heterozygous for mutation of the *Hoxc9,10,11* genes were almost completely infertile. In this report we show that *Hoxc9,10,11* genes have redundant function with *Hoxa9,10,11* and *Hoxd9,10,11* in uterine gland formation.

Single cell RNA-seq (scRNA-seq) is a powerful tool for the dissection of normal and mutant development (Potter, 2018). It can define the global gene expression states of the multiple cell types present in a developing organ. Analysis using scRNA-seq can help determine how early lineage decisions are made (Brunskill et al., 2014). It can characterize the transcription factor codes that define different cell types, and it can provide a comprehensive analysis of potential ligand-receptor interactions. In this report we used scRNA-seq to examine the wild type developing uterus at PND12, as early gland formation is taking place. In addition, we used scRNA-seq to examine the perturbed gene expression patterns of all cell types of the PND12 *Hoxa9,10,11*+/-,*Hoxc9,10,11*+/-, *Hoxd9,10,11*+/- (ACD+/-) mutant uteri, with heterozygous mutation of 9 Hox genes, as well as *Hoxa9,10,*+/-,*Hoxc9,10,11*+/-, *Hoxd9,10,11*+/- (ACD+/- WTA11), with two wild type *Hoxa11* genes and heterozygous mutation of 8 Hox genes. The results create a single cell resolution atlas of the gene expression patterns of the wild type developing uterus and also define the changed gene expression profiles of all cell types in mutant uteri. We observed a compelling disruption of Wnt signaling in the Hox mutants, as well as strongly reduced stromal expression of *Cxcl12* (SDF1), which encodes a ligand for Cxcr4, expressed by the LE. In multiple developing contexts Cxcr4/Cxcl12 interaction has been previously shown to drive cell migration and tubulogenesis (Doitsidou et al., 2002; Feng et al., 2014; Ivins et al., 2015; Molyneaux et al., 2003; Ueland et al., 2009), key steps in uterine gland formation.

## Results

### ACD+/- female mice are sub-fertile

The roles of *Hoxa9,10,11* and *Hoxd9,10,11* in male and female fertility have been previously examined (Raines et al., 2013). A modified recombineering method was used to introduce simultaneous frameshift mutations in the flanking sets of Hox genes on the HoxA and HoxD clusters (Raines et al., 2013). Both male and female *Hoxa9,10,11-/-* mice are infertile, consistent with earlier studies of *Hoxa10-/-* and *Hoxa11-/-* mice (Gendron et al., 1997; Satokata et al., 1995). *Hoxa9,10,11*+/- heterozygous females gave normal litter sizes while *Hoxd9,10,11*-/- homozygous mutant females showed only a slight reduction of litter size (8.7+1.3, average + <u>s.e.m.</u> vs WT of 11.5+1.8), but combined mutant *Hoxa9,10,11*+/-, *Hoxd9,10,11-/-* females were completely infertile, with vaginal plugs never producing litters (Raines et al., 2013). The synergistic severity of phenotype shows a function for the *Hoxd9,10,11* genes that is redundant with the *Hoxa9,10,11* genes.

*Hoxc10* and *Hoxc11* are also expressed in the human female uterus, suggesting possible fertility function (Akbas and Taylor, 2004). Frame shift mutations were simultaneously introduced into the first exons of *Hoxc9, Hoxc10*, and *Hoxc11* using recombineering (Drake et al., 2018). It is important to note that this approach leaves shared enhancer regions intact, thereby reducing impact on the expression of the remaining Hox genes not mutated. Female *Hoxc9,10,11-/-* mice mated to wild-type males had similar size litters (9.33±0.88 pups/litter, n =3) to those from wild-type matings (9.92±0.35, n = 13). To further test for possible *Hoxc9,10,11* fertility function we then used a “sensitized” mouse that was heterozygous for the six *Hoxa9,10,11*+/- *Hoxd9,10,11+/-* (AD+/-) genes, which results in reduced fertility with 5.3 pups per litter (Raines et al., 2013). When we made mice heterozygous for all nine *Hoxa9,10,11, Hoxc9,10,11, Hoxd9,10,11* (ACD+/-) genes we observed a dramatic decrease in fertility. From a total of 41 vaginal plugs only two litters born, with a total of 6 pups, giving about 0.15 pups per vaginal plug. The further reduction of fertility in the ACD+/- mice compared to AD+/- gives evidence for fertility function for the *Hoxc9,10,11* genes.

The fertility functions of the *Hoxc9,10,11* genes were also shown through crosses with only the *Hoxd9,10,11* mutants. While both the *Hoxc9,10,11*-/- and the *Hoxd9,10,11*-/- mice gave very near wild type litter sizes, the combined mutant *Hoxc9,10,11*+/-, *Hoxd9,10,11*-/- mice gave an average litter of only 5.4 pups (18 litters with a total of 97 pups), while the converse *Hoxc9,10,11*-/-, *Hoxd9,10,11*+/- mutants gave an average litter of only 5.8 pups (16 litters with a total of 92 pups). This again gives evidence for *Hoxc9,10,11* fertility function that was previously concealed by functional redundancy with other paralogous Hox genes.

We also generated ACD+/- mice but with two wild type *Hoxa11* alleles (*Hoxa9,10*^*+/-*^, *c9,10,11*^*+/-*^, *d9,10,11*^*+/-*^)(ACD+/-WTA11). As noted previously, *Hoxa11* is known to play a critical role in both male and female fertility (Gendron et al., 1997; Hsieh-Li et al., 1995; Lewis et al., 2003). Four *Hoxa9,10*^*+/-*^, *c9,10,11*^*+/-*^, *d9,1011*^*+/-*^ females were mated with wild type males, giving twelve litters totaling 125 pups, or 10.41±0.77 pups per litter, which is comparable to wild type. It is remarkable that the addition of one wild type *Hoxa11* allele to the ACD+/- mice, with nine mutant Hox alleles, can rescue a wild type phenotype. This shows the relative importance of *Hoxa11* in female fertility.

In this study we focus on the ACD+/- genotype, which shows a highly penetrant female infertility, yet the mice are quite healthy, without the severe kidney malformations associated with removal of additional Hox10,11 genes.

### ACD+/- mice show a dramatic reduction in the number of uterine glands

Histologic analysis of the ACD+/- uteri showed a reduction in the number of uterine glands (Fig. 1D-F) compared to WT uteri (Fig. 1A-C). At PND10, PND20 and adult the luminal epithelium (LE), mesenchymal stroma, and the muscle layers of ACD+/- mice appeared normal (Fig. 1). At all stages examined, however, there was a dramatic reduction in the number of uterine glands. The most striking difference was seen in the adult uterus (Fig. 1). Glands were counted from transverse sections of uteri. Wild-type adult uteri contained 32.63 ± 2.46 glands/section (n = 13), while ACD+/- uteri contained 3.44±0.66 glands/section (n = 7). These results give evidence for the sub-fertility being the result of a drastic reduction in the number of uterine glands, which supply vital proteins, including Lif (Rosario et al., 2014) and calcitonin (Zhu et al., 1998), which are required for embryo implantation. Interestingly, the adult ACDWTA11+/- mice, with two wild type *Hoxa11* alleles, had wild type numbers of uterine glands (37.0+/-6.66 glands/section, n = 3), again showing the importance of normal *Hoxa11* expression in female fertility.

**Figure 1.**
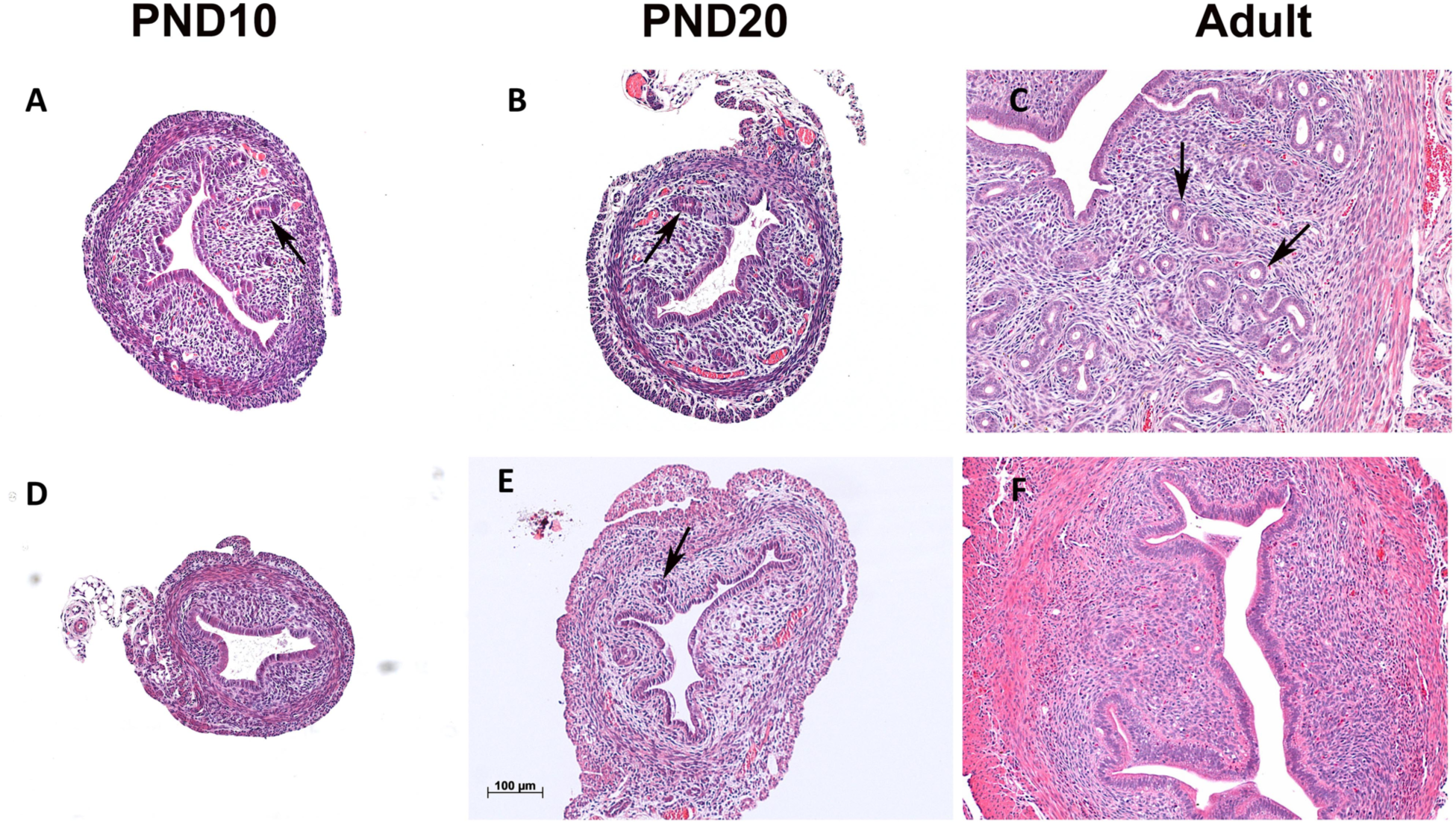
Reduced gland formation in ACD+/- Hox mutant uteri. At PND10, PND20 and adult the ACD+/- mutant uteri (D,E,F) show fewer glands than wild type (A,B,C). Arrows mark selected glands. H&E stains.

The ACD+/- mice produce normal numbers of embryos. WT and ACD+/- mutant females were mated with WT males. The detection of a vaginal plug equaled gestational day 1 (GD1). Mice were euthanized at GD4 and one horn of each uterus was gently flushed. Similar numbers of blastocysts were isolated from wild-type (4.45+/-0.72 blastocysts/uterine horn) and ACD+/- mice (3.38+/-0.54 blastocysts/uterine horn). Recovered blastocysts from ACD+/- mice were morphologically normal (data not shown).

### Single cell RNA-Seq analysis of the wild type and ACD+/- and ACD+/-WTA11 mutant developing uteri

scRNA-seq is a powerful technology that can give a high resolution definition of the gene expression programs that drive development (Adam et al., 2017; Brunskill et al., 2014; Camp et al., 2017; Magella et al., 2018; Potter, 2018; Treutlein et al., 2014). To better understand the normal and perturbed gene expression patterns of the developing uterus we carried out scRNA-seq on the WT, ACD+/-, and ACD+/-WTA11 uteri at PND12. We were particularly interested in uterine gland formation and how this process might be disrupted in the ACD+/- mutants. Gland formation begins with the budding of the luminal epithelium at approximately PND6 and continues into adulthood during endometrial regeneration following menstruation in humans and following birth in both humans and mice (Bigsby et al., 1990; Brody and Cunha, 1989; Cooke et al., 2012; Garry et al., 2010; Huang et al., 2012).

### Wild Type

scRNA-Seq was performed using Drop-Seq (Macosko et al., 2015). A total of 6,343 WT cells passed quality control filters. Unsupervised clustering of cells was carried out using the Seurat R package as previously described (Butler et al., 2018). A t-SNE plot showing the resulting clusters is shown in Fig. 2A. Major cell types were defined using the expression of known cell type-specific genes, with epithelial cells (n=1487) expressing *Epcam* and *Cdh1*, endothelial cells (n=373) expressing *Pecam1* and *Emcn*, stromal cells (n=2241) expressing *Pdgfra, Dcn,* and *Col15a1*, myocytes (n=1749) expressing *Mef2a* and *Pdlim3*, lymphocytes (n=106) expressing *Cd52*, and *Ptprc*, myeloid cells (n=39) expressing *Lyz2, Pf4*, and *C1qc,* pericyte**s** (n=269) expressing *Rgs5* and *Cspg4*, and mesothelial cells (n=79) expressing *Upk3b* and *Lrrn4*. A heatmap of the top 15 marker genes from each cluster is shown in Fig. 2B. Lists of genes with compartment enriched expression are presented in Table S1.

**Figure 2.**
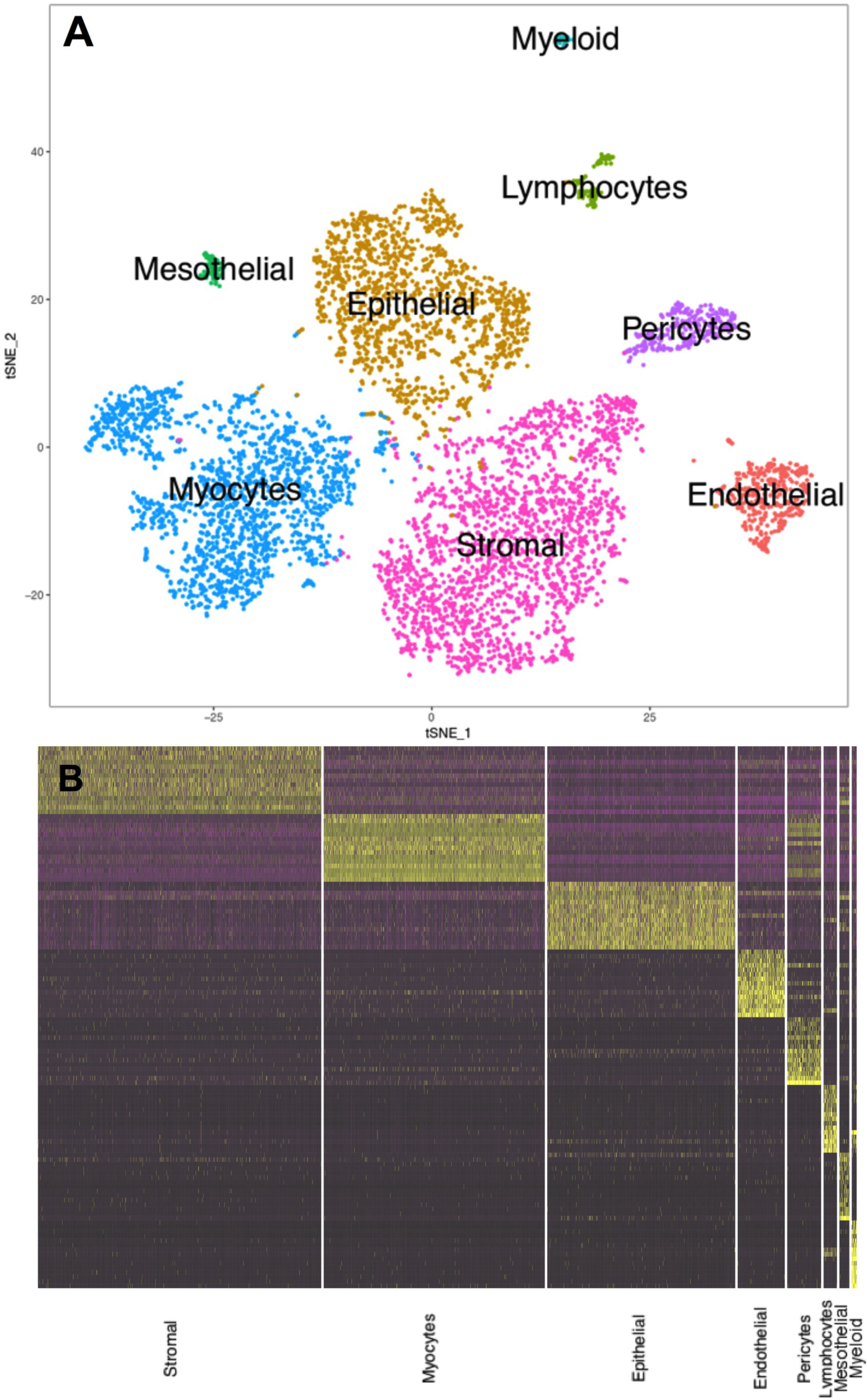
Single cell RNA-seq analysis of the developing uterus. (A) The t-SNE plot shows unsupervised clustering of wild type PND12 cells into eight groups, as labeled. (B) The heatmap shows differential expression of the top fifteen genes for each cell type, with yellow indicating high relative gene expression, purple indicating low expression, and black indicating no expression.

### Subclusters

Five of the eight cell types were then separated and subclustered (Fig. 3A-F). Elevated expression of markers of sub-compartments is shown in the t-SNE plots of Fig. 3. Using the Split Dot Blot function of the Seurat package, a gene expression profile of each sub-cluster was generated (Fig. 4A). The size of each circle represents the percentage of cells within a cluster expressing the gene of interest. For example, both wild-type luminal epithelial (LE_WT) and germinal epithelial (GE_WT) cells express *Ifitm1.* These two cell populations can be differentiated by the higher percentage of cells expressing *Calb1* in LE_WT in comparison to GE_WT (Fig. 4A). A heatmap depicting the top 15 marker genes for each sub-cluster shown in Fig. 4B. The complete lists of genes with elevated expression in each sub-cluster are presented in Tables S2.

**Figure 3.**
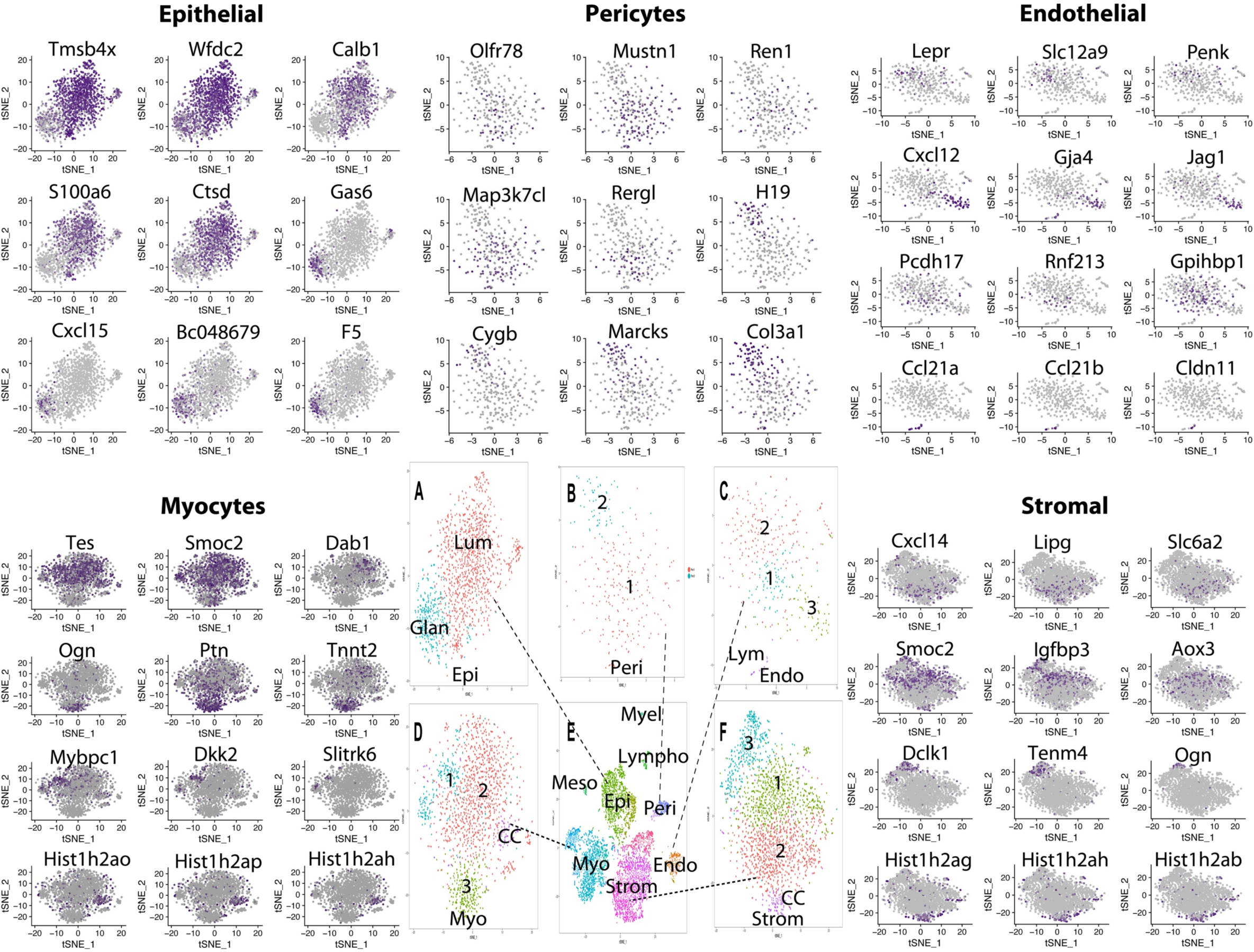
Defining cell subtypes. Five cell types were subjected to subclustering. Expression patterns of selected genes associated with cell subtypes are shown in surrounding t-SNE plots. (A) Epithelial cells divided into glandular (*Gas6, Cxcl15, Bc048679, F5*) and luminal (*Tmsb4x, Wfdc2, Calb1, S100a6, Ctsd*), (B) pericytes divided into two subtypes, type 1 (*Olfr78, Mustn1, Ren1, Map3k7cl, Rergl*) and type 2 (*H19, Cygp, Marcks and Col1a1*), (C) endothelial cells divided into 4 subtypes including lymphatic, expressing *Ccl21a, Ccl21b* and *Cldn11*, type 1 (capillary) expressing *Pcdh17, Rnf213 and Gpihbp1*, type 2 (venous) expressing *Lepr, Slc12a9 and Penk* and type 3 (arterial) expressing *Cxcl12, Gja4 and Jag1*. (D) Myocytes and (F) stromal cells each dividing into four subtypes as shown.

**Figure 4.**
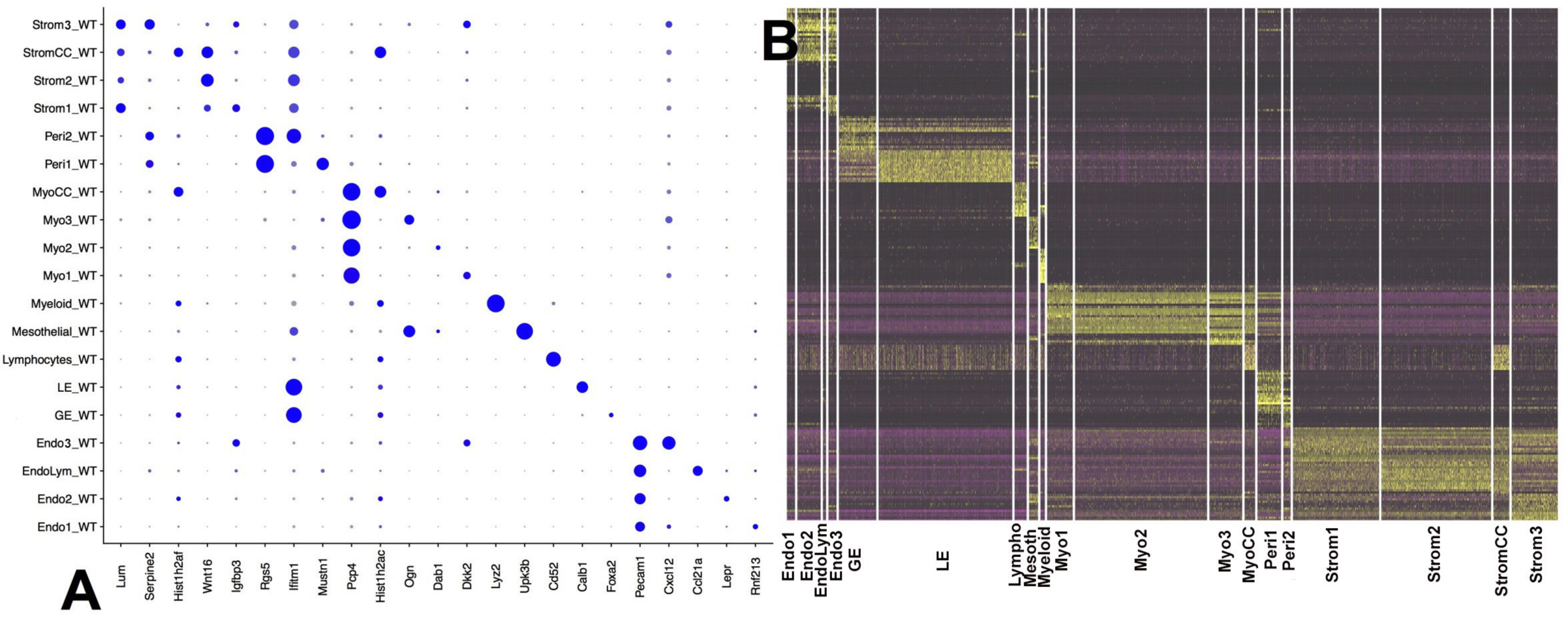
Gene expression patterns of cell types and subtypes. (A) Split Dot Blot shows cell types and selected differentiating genes, with the size of the circle representing the percentage of cells showing expression. (B) Heatmap showing gene expression relationships of cells types.

Genes with elevated expression in GE versus LE included *Foxa2, Wif1, Gas6, Ccnd1, Cxcl15, BC048679, F5*, and *Egfl6*, which were previously shown to have elevated GE expression in a microarray study (Filant and Spencer, 2013). Genes more strongly expressed in LE cells compared to GE included *Tmsb4x, Wfdc2, 1600014C10Rik, Calb1, S100a6* and *Ctsd*. Mesothelial cells (n = 79) are specialized squamous cells that form a monolayer that lines the body’s serous cavities and internal organs, providing a slippery, non-adhesive and protective surface. Genes with elevated expression in mesothelial cells included *Upk1b, Upk3b, Cldn15, Fgf1*, and *Fgf9*.

Four stromal cell subtypes were identified on the basis of unsupervised subclustering (Fig. 4). All four stromal populations expressed *Lum, Igfbp4, Pdgfra*, and *Col15a1*, while subclusters were distinguished by expression of *Smoc2, Igfbp3, Aox3,* and *Gem* (Stromal 1), *Cxcl14, Lipg, 2700046A07Rik*, and *Slc6a2* (Stromal 2), *Dclk1, Tenm4, Ogn,* and *C1qtnf3* (Stromal 3)(Fig. 4). A fourth subcluster was marked by high expression of cell proliferation, cell cycle genes (StromalCC). The scRNA-seq data therefore reveals an interesting diversity of stromal cell subtypes.

In similar fashion, four myocyte subtypes were identified, with all expressing the muscle related genes *Mef2a, Myl6, Myh11*, and *Tpm1*, and subclusters separated by differential expression of *Mybpc1* [in striated muscle bands], *Dkk2, Slitrk6,* and *Dio2* (Myo1), *Tes, Smoc2* [smooth muscle associated protein], *Dab1,* and *Camk1d* (Myo2), *Ogn, Ptn, Tnnt2*, and *Ccnd,* (Myo3) and a fourth subcluster (MyoCC) with elevated expression of cell cycle genes and proliferation associated genes, including *Top2a, Cdk1, Ccnb2, Histih2ao, Hist1h2ap, Hist1hadh*, and *Hist1h2an*. Of interest, the Myo1 group has some skeletal muscle properties, with Mybpc1 found in c bands of skeletal muscle and Dkk2 showing strongest expression in skeletal muscle, while the Myo2 cells express genes associated with smooth muscle (Smoc2 and Tes).

Two pericyte sub-clusters were also identified, both expressing the known pericyte markers *Notch3, Rgs5, Cspg4*, and *Pdgfrb* (He et al., 2016; Wang et al., 2014). Pericyte sub-group 1 cells also expressed, *Mustn1, Ren1, Tagln, Map3k7cl,* and *Rergl* while Pericyte sub-group 2 cells expressed *H19, Cygb, Marcks, Col3a1, as well as Ifitm1* and *Art3,* membrane proteins associated with pericytes (He et al., 2016). Of interest, *Marcks, Col3a1* and *Ifitm1* all show elevated expression in brain microvascular cells (Daneman et al., 2010). *Cygb* encodes a pericyte globin required to prevent premature senescence and age dependent malformations of multiple organs (Thuy le et al., 2016).

Four endothelial cell sub-clusters were identified with all clusters expressing the panendothelial markers *Pecam1* and *Emcn*. The vascular endothelial markers *Cd34* and *Sox17* were expressed in every sub-cluster except EndoLym (lymphatic). The Endo1 cell cluster expressed *Pcdh17, Rnf213, Sik3,* as well as *Gpihbp1,* which is expressed exclusively in capillaries (Young et al., 2011). The Endo2 cell cluster showed elevated expression of *Nrp2, Lepr, Slc16a9, Penk, and Fendrr,* indicating venous cell identity (dela Paz and D’Amore, 2009). As expected the EndoLym cell cluster expressed *Prox1* and *Lyve1* as well as *Ccl21a, Ccl21b, Cldn11,* and *Fxyd6,* while the Endo3 cells expressed *Cxcl12, Gja4, Jag1,* and *Dkk2.* Both *Gja4* (Villa et al., 2001), and *Jag1* (Fang et al., 2017) expression have been linked to endothelial arterial cell specific differentiation.

### Analysis of growth factor-receptor interactions

The scRNA-seq data provides a global view of the gene expression patterns of the multiple cell types of the developing uterus. It defines the transcription factor codes that drive the distinct lineage differentiation programs. Further, it describes the combinations of growth factors and growth factor receptors expressed by each cell type. We were particularly interested in the growth factor/receptor interactions taking place between the stromal cells and luminal epithelial cells that might play a role in promoting uterine endometrial gland formation.

Most of the Hox genes targeted in the ACD+/- mutants showed strongest expression in the stromal cells at PND12, with none showing significant expression in the epithelial compartment (Fig. 5). Indeed, for the HoxA cluster only the *Hoxa9,10,11* genes showed significant expression in the developing PND12 uterus, with each expressed in the stromal and myocyte compartments (Fig. 5). For the HoxB cluster none of the genes showed strong expression in stromal cells, but many showed significant expression in the epithelial and endothelial compartments. Of particular interest, none of the HoxC cluster genes showed strong expression in the developing uterus, although *Hoxc9, Hoxc10* and *Hoxc11* were upregulated in the ACD+/- mutant uterus (see below). For the HoxD cluster the *Hoxd3,8,9,10,11* genes showed expression in the PND12 uterine stromal, myocyte and endothelial compartments (Fig. 5).

**Figure 5.**
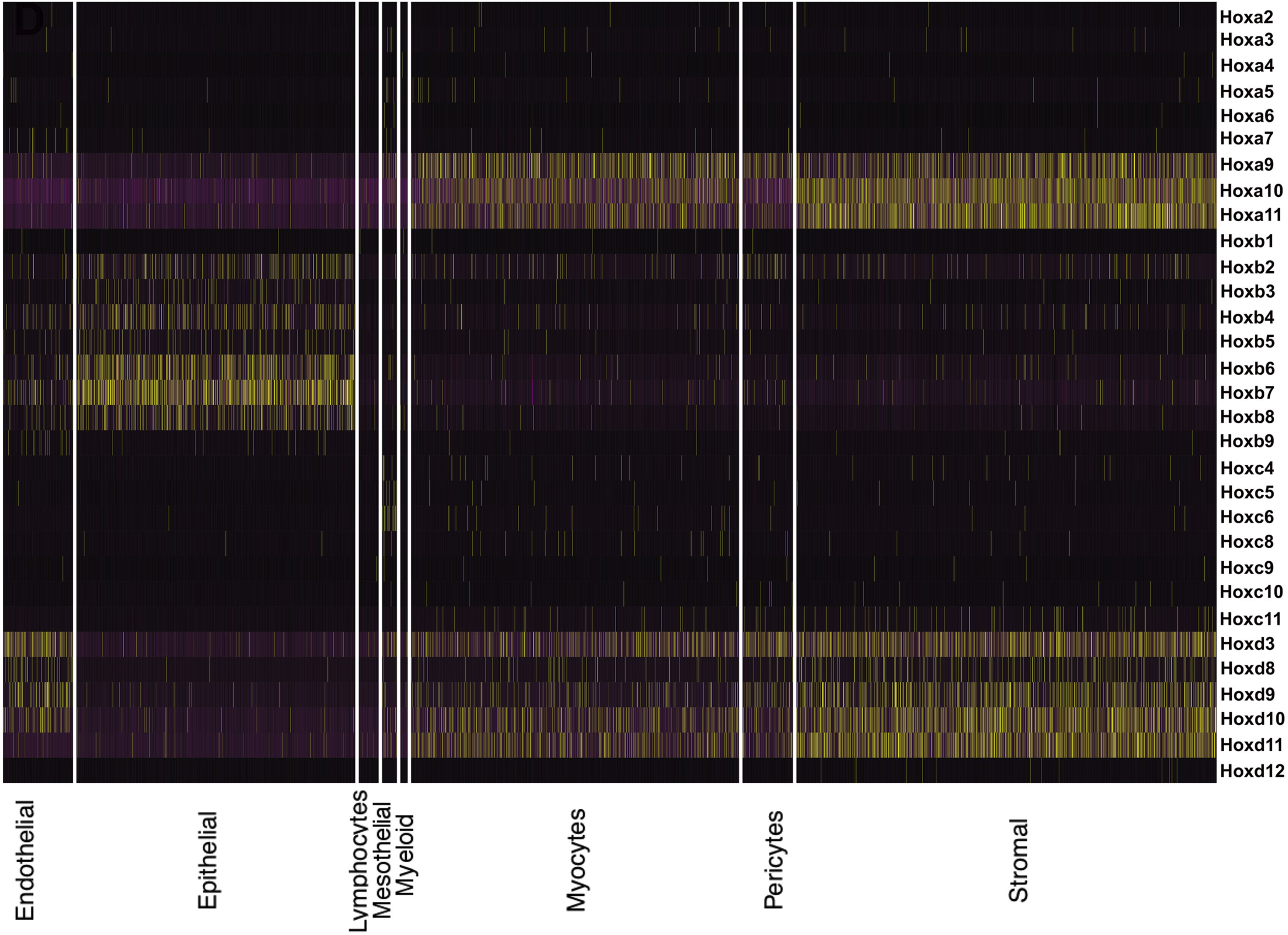
Expression of Hox genes in the PND12 developing wild type uterus. For the HoxA cluster only *Hox 9,10,11* genes show significant expression, with all three showing very similar expression levels in the stromal, myocyte and pericyte cells. HoxB cluster gene expression is mostly restricted to epithelial cells. The HoxC cluster shows very little expression in the developing wild type uterus, although there is compensatory upregulation in the ACD+/- uterus (Fig. 9). *Hoxd3,9,10,11* are expressed in the stromal, pericyte, myocyte and endothelial cell types. Only Hox genes with detected expression are shown.

This stromal expression of the ACD+/- mutated Hox genes, and absence of their expression in LE, suggests that the reduced invagination of LE cells to form glands in the ACD+/- mutants could be the result of altered cross talk between stroma and LE. Therefore, we restricted the analysis to growth factors expressed in stromal cells and their corresponding receptors in any cell type. Growth factors and receptors were filtered requiring detected expression in greater than 7.5% of cells in a cluster. A Circos plot was generated that displays resulting potential growth factor/receptor interactions (Fig. 6)(Krzywinski et al., 2009). Of particular interest, stromal cells were found to express *Bmp, Cxcl12, Gdf10, Igf1, Manf, Rspo3* and *Tgfbeta*, all potentially able to bind to their corresponding receptors on epithelial cells.

**Figure 6.**
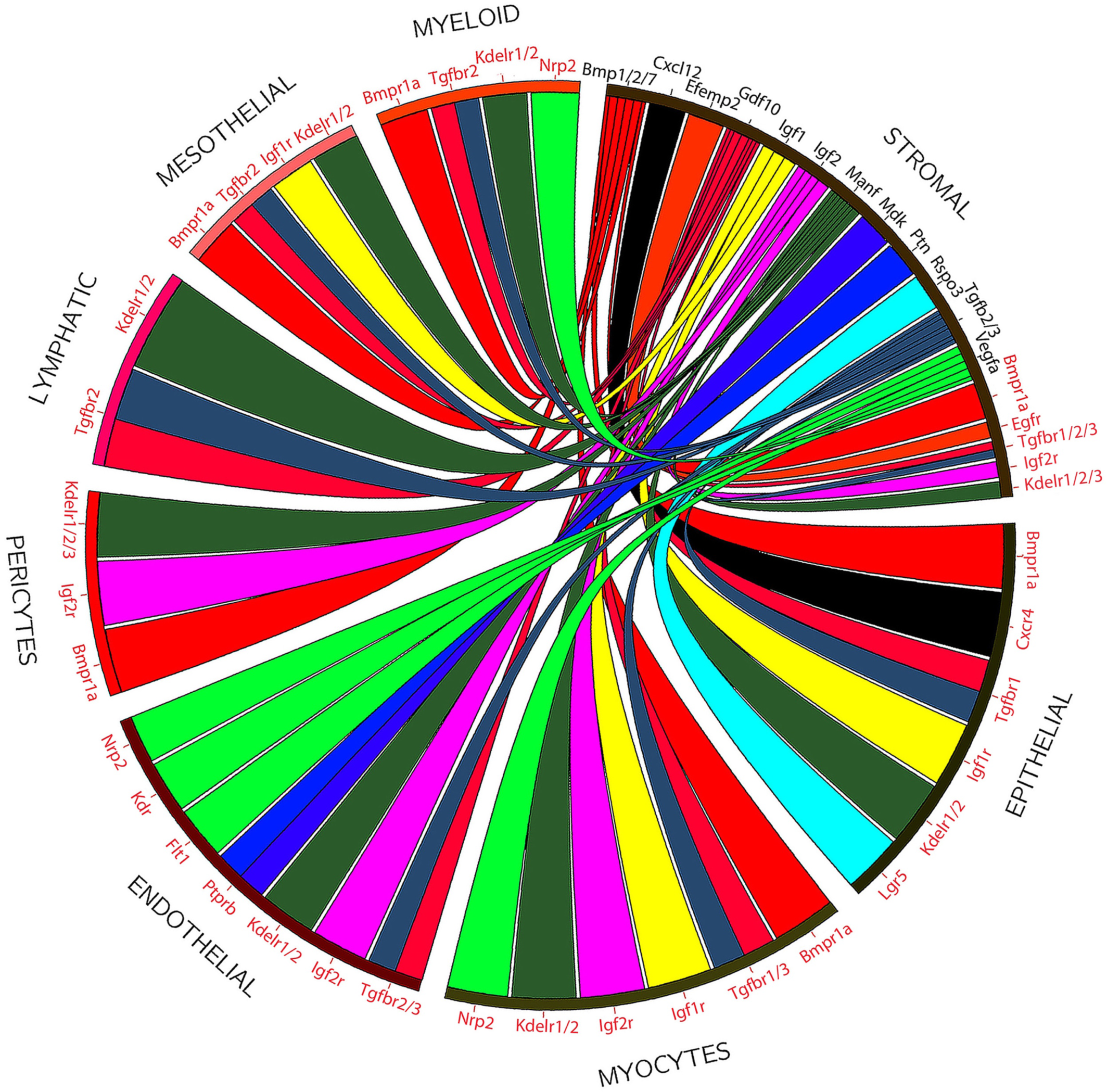
Potential ligand receptor interactions in the developing uterus. Circos plot showing ligands made by stromal cells (black) and receptors made by various cell types (Dastani et al.), and potential interactions. The lists are not all inclusive due to filtering used to restrict the number of interactions.

The stromal growth factors and their corresponding epithelial receptors were then used to generate a network plot using GeneMania software (Warde-Farley et al., 2010) (Fig. 7). The functional GO enrichment analysis indicates that some of the genes within the network are involved in gland development (3.98E-16), morphogenesis of branching epithelium (3.54E-16), morphogenesis of a branching structure (7.67E-16), branching morphogenesis of an epithelial tube (1.46E-15), and epithelial tube morphogenesis (3.21E-12), consistent with a role in the initiation and/or progression of uterine gland development.

**Figure 7.**
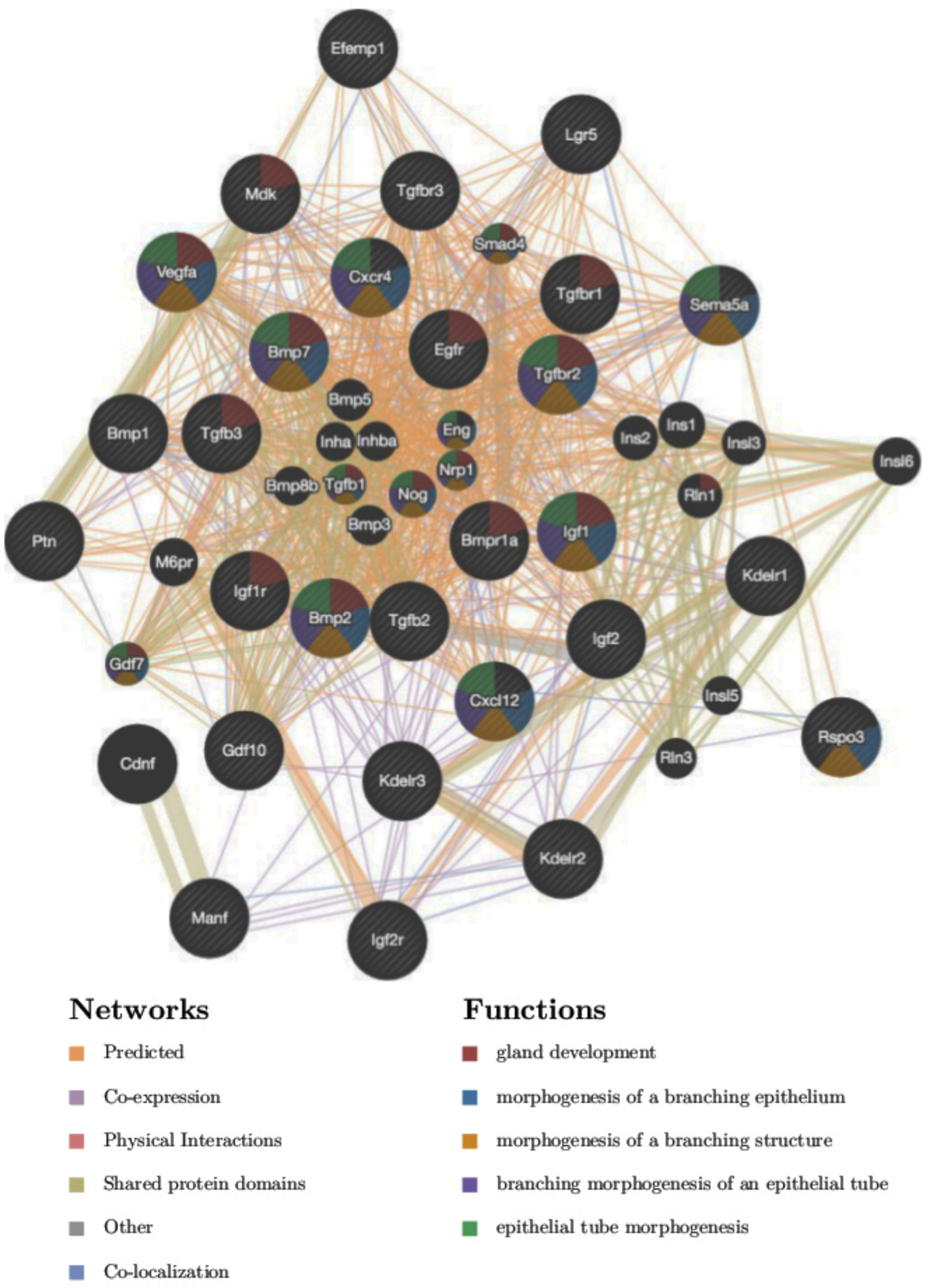
Functional analysis of expressed stromal ligand/epithelial receptors. The network plot was made using GeneMania software (Warde-Farley et al., 2010). The functional GO enrichment analysis indicates that some of the genes within the network are involved in gland development (3.98E-16), morphogenesis of branching epithelium (3.54E-16), morphogenesis of a branching structure (7.67E-16), branching morphogenesis of an epithelial tube (1.46E-15), and epithelial tube morphogenesis (3.21E-12), consistent with a role in the initiation and/or progression of uterine gland development.

### scRNA-Seq analysis of WT, ACD+/- and ACD+/-WTA11 uterine cells

To better understand the altered gene expression patterns in each mutant cell type we also carried out scRNA-seq for ACD+/- and ACD+/-WTA11 PND12 uteri. All cells of the three genotypes were combined and clustered in an unsupervised manner resulting in 11 major cell types (Fig. 8, right panel). The clustered cells could then be color coded according to genotype (Fig. 8, left panel). Of interest, there were striking genotype specific differences for three clusters, the stromal, myocyte and epithelial cells. For example, within the stromal cluster the cells of the three genotypes clearly subclustered separately. The WT and ACD+/- cells were at the two ends of the cluster, with the ACD+/-WTA11 cells in between. For the myocyte cluster there was a similar result, with the WT and ACD+/- cells at the two ends of the cluster and once again the ACD+/-WTA11 cells in between. For the epithelial cell cluster there was also genotype dependent cell separation, although not as complete as observed in the stromal and myocyte clusters.

**Figure 8.**
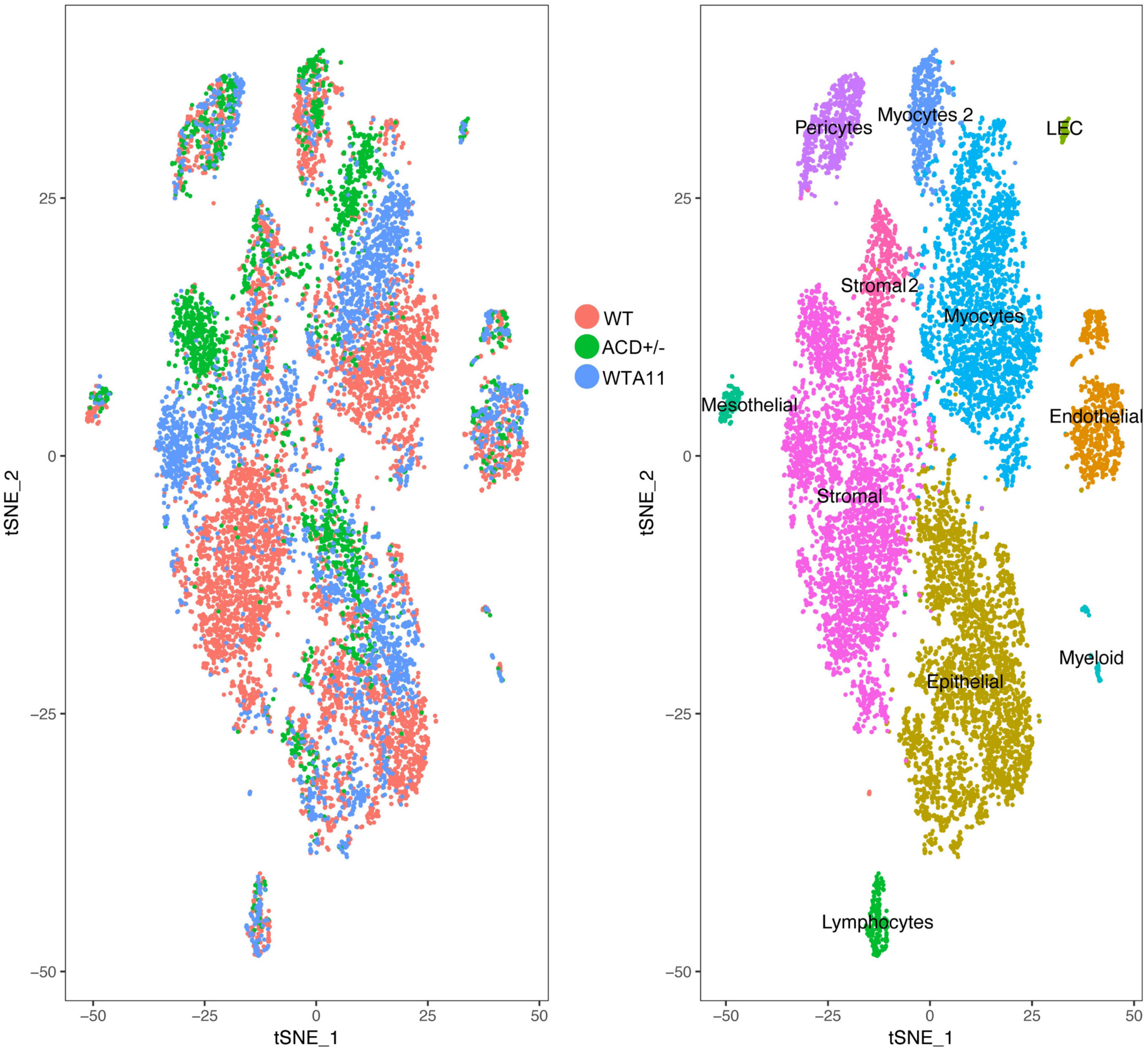
Comparison of WT, ACD+/- and ACD+/-WTA11 cells. Cells of all three genotypes were combined for unsupervised clustering analysis. Right panel shows t-SNE plot color coded according to resulting cells types, as labeled. Left panel shows the same t-SNE only color coded according to genotype. Of interest, the two cells types, stromal and myocyte, with strongest expression of the mutated Hox genes show very distinct cell separations based on genotype. For example, the stromal cluster includes ACD+/- cells grouping at the top, WT cells at the bottom, and ACD+/-WTA11 cells in between. Myocytes show a very similar separation. Other cell types, such as endothelial, lymphocytes and pericytes, with little expression of the Hox genes mutated, show no such separation based on genotype, arguing against a simple batch effect explanation.

We then examined the differential expression of WT versus ACD+/- stromal cells to find possible changes in gene expression that might result in perturbed stromal-LE signaling. Genes involved in signaling with reduced expression in the mutant stroma included *Thbs1, Thy1, Igfbp6, Gas6, Cd44, Ndp*, and *Cxcl12*. Thbs1 is a major activator of Tgf-beta (Crawford et al., 1998), so reduced *Thbs1* expression in the mutant stroma suggests reduced Tgf-beta signaling. As noted earlier, Tgf-beta is expressed by stromal cells and its receptor is expressed by LE. Also of interest, Thy1 can interact with integrins to drive cell adhesion/migration responses (Barker and Hagood, 2009), which could play a role in GE formation. *Gas6* mutant mice are viable and give normal size litters (Yanagita et al., 2002), arguing against a key role for *Gas6* in GE formation.

Of particular interest, the *Ndp* gene with reduced expression in mutant stroma encodes Norrin protein, which can interact with frizzled receptor to drive canonical Wnt signaling (Zhang et al., 2017), which in turn is critically important for GE formation. Consistent with reduced Wnt signaling in the mutant there was increased expression of *Dkk2* (1.9 FC), which encodes a Wnt inhibitor. The mutants, however, also showed increased expression of *Rspo1* and *Rspo3*, which activate canonical Wnt signaling. We also observed decreased expression of *Gli1* and *Ptch2* in mutants, suggesting reduced Hedgehog signaling.

Finally, *Cxcl12*, also known as stromal cell derived factor 1, was strongly down regulated 3.8 fold change (FC) in the ACD+/- mutant stroma. Of interest, Cxcl12 drives angiogenesis, tubulogenesis and cell migration (Belmadani et al., 2005; Molyneaux et al., 2003; Pi et al., 2009; Ueland et al., 2009; Zhang et al., 2005) through interaction with its Cxcr4 receptor, which is expressed in the LE.

### RT-PCR analysis of gene expression in mutant uteri

We carried out RT-PCR to further validate selected gene expression differences in mutants identified through scRNA-seq. First, we were interested in the altered expression of the *Hoxc9,10,11* genes themselves, which showed low expression in the developing uterus (Fig. 6). RT-PCR showed that expression of these genes is upregulated in the ACD+/- mutant uterus (Fig. 9). These genes show upregulation even though each is heterozygous for a frameshift mutation in the first exon that would be expected to result in increased nonsense mediated decay. These results suggest that the *Hoxc9,10,11* genes normally play a limited role in early uterus development but undergo increased compensatory expression when the dose of *Hoxa9,10,11* and *Hoxd9,10,11* genes is reduced.

**Figure 9.**
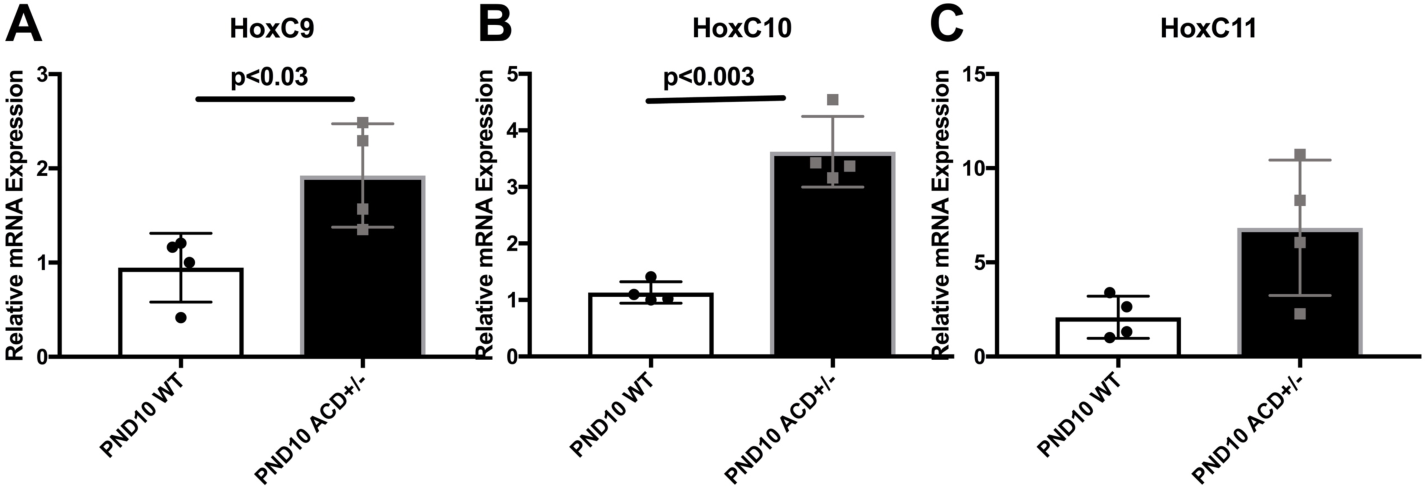
Compensatory expression of *Hoxc9,10,11* genes in PND10 ACD+/- mutant uteri. RT-PCR was used to compare expression levels of *Hoxc9,10,11* in WT and ACD+/- mutant uteri at PND10. There appeared to be significant upregulation in ACD+/- mutants, even though all of the *Hoxc9,10,11* genes were heterozygous mutant, carrying a first exon deletion resulting in a frameshift mutation that that should result in nonsense mediated decay for half of the transcripts.

RT-PCR was used to confirm the reduced expression of *Lef1, Sox9, Sp5, Gli1, Ptch2*, and *Cxcl15* and in ACD+/- mutant uteri compared to WT. We examined both WT and ACD+/- PND10 and PND20 uteri. The Lef1 transcription factor is a key mediator of Wnt signaling but is also a key target, with Wnt signaling reporter transgenes often including concatemerized Lef binding sites (Barolo, 2006). The observed significant reduction of *Lef1* expression therefore gives evidence for reduced Wnt signaling in the ACD+/- mutant uterus (Fig. 10). The observed reduced expression of *Sp5*, a known Wnt target (Huggins et al., 2017), is also consistent with reduced Wnt signaling (Fig. 10).

**Figure 10.**
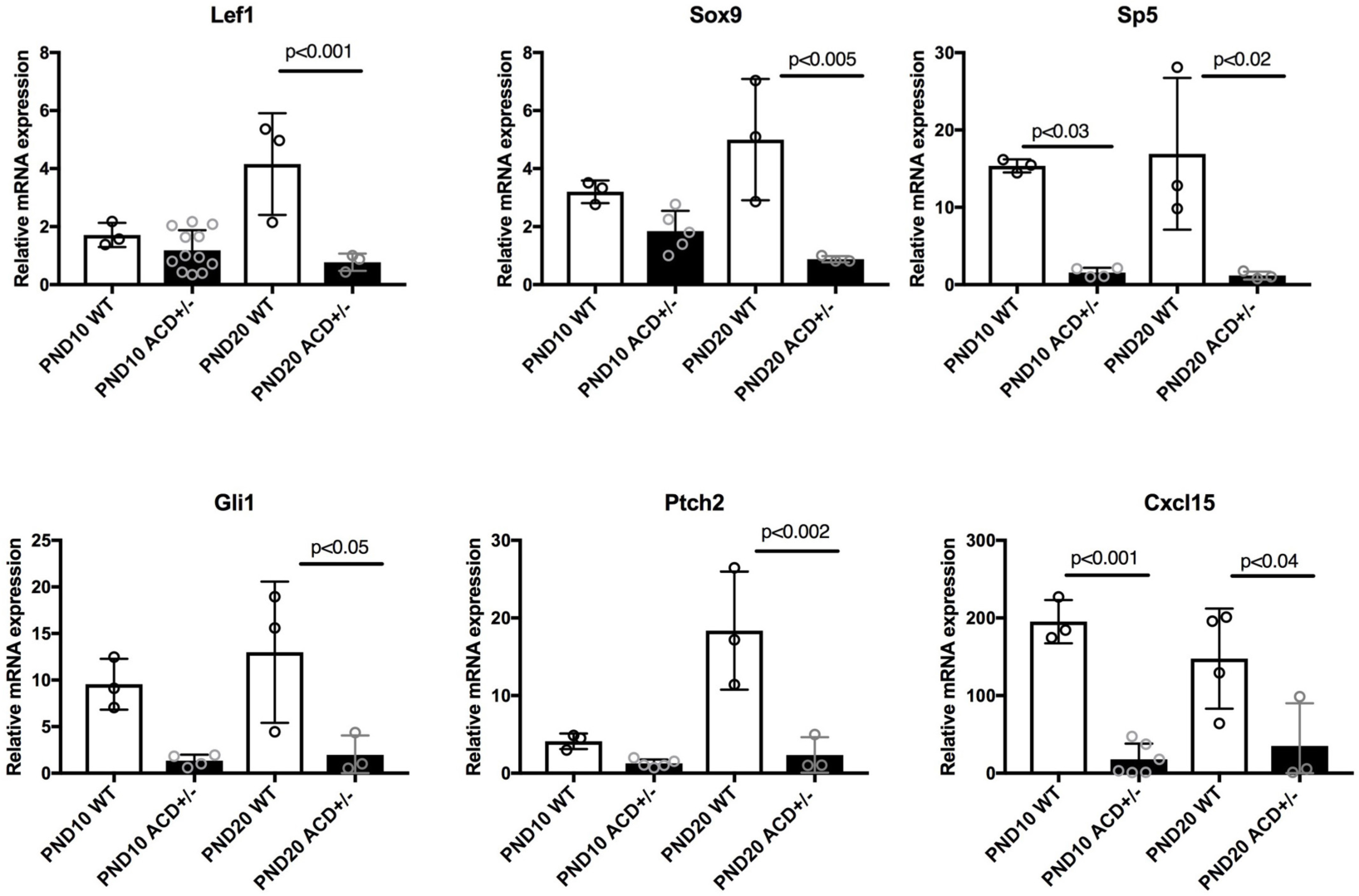
RT-PCR analysis of altered gene expression in ACD+/-mutants. RNA was isolated from PND10 and PND20 wild type and mutant whole uteri. There was reduced mutant expression of *Lef1*, consistent with reduced Wnt signaling in mutants. *Sp5*, and target of Wnt signaling, also showed reduced expression in mutants. *Gli1* and *Ptch2* also showed lower expression in mutants, suggesting perturbed hedgehog signaling. *Cxcl15* and *Sox9* have highest expression in the GE, and their reduced expression therefore likely reflects reduced GE representation in mutants.

The scRNA-seq and RT-PCR data also provide evidence for reduced hedgehog signaling. *Gli1* and *Ptch2* encode integral components of the hedgehog signaling pathway and are also targets of hedgehog signaling (Hooper and Scott, 2005). Expression of Gli and Ptch are increased by hedgehog signaling and thus they serve of markers of hedgehog signaling (Chen and Struhl, 1996; Marigo et al., 1996). The RT-PCR validation of reduced expression of both *Gli1* and *Ptch2* in the mutant uterus therefore gives evidence for decreased hedgehog signaling (Fig. 10). In addition, RT-PCR showed lowered expression levels for *Sox9* and *Cxcl15*. The *Cxcl15* gene shows restricted expression in the GE of the uterus (Table S1)(Kelleher et al., 2016), and its reduced expression in ACD+/- uteri is therefore likely a simple reflection of reduced GE representation in the mutants.

### Immunhistochemistry

We further examined the reduced Wnt signaling in ACD+/- uteri using immunostaining with Lef1 antibody. At PND10 there was dramatic reduction of Lef1 expression in ACD+/- mutants (Fig. 11, top panels). We then asked if this reduced expression persisted in adults. We observed that during diestrus, estrus, and also in the E5 post vaginal plug uterus there was significant reduction of Lef1 expression in mutants (Fig. 11). In WT adult uteri Lef1 expression was restricted to stroma during diestrus and at E5, but during estrus Lef1 was expressed in both stroma and GE. Lef1 was also expressed in budding LE that was forming GE at PND10. In mutants the very low levels of Lef1 expression observed appeared restricted to stroma.

**Figure 11.**
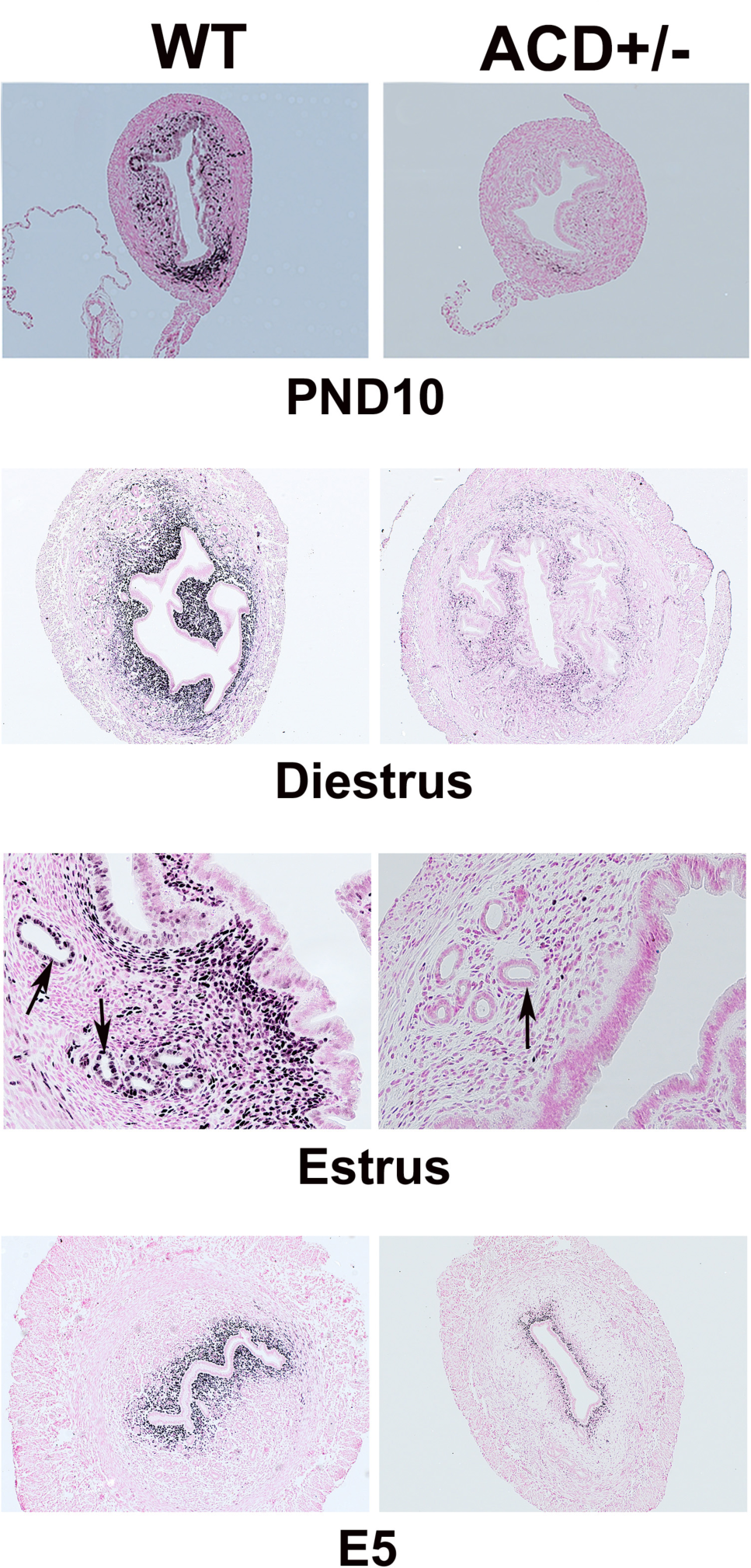
Immunohistochemical analysis of Lef1 expression. At PND10 there was robust Lef1 expression in uterine stromal cells, in particular at the mesometrial pole. In ACD+/- mutants (right panels) Lef1 expression was greatly reduced. Reduced Lef1 expression in mutants was also seen in adults at diestrus, estrus and at E5 post fertilization. During estrus there was Lef 1 expression in GE as well as stroma in WT uterus. Arrows point to uterine glands.

## Discussion

In this study we identify novel female fertility function for the *Hoxc9,10,11* genes, with ACD+/- mice showing a much more severe infertility phenotype than AD+/- mice. Histologic analysis showed an approximate ten fold reduction in the number of uterine glands in ACD+/- mice compared to wild type, giving evidence for a role for *Hoxc9,10,11* genes in the formation of uterine glands, which are required for fertility.

Single cell RNA-seq was then used to explore the gene expression programs driving distinct lineage trajectories in the wild type developing uterus, as well as to define the perturbed gene expression patterns in the multiple cell types of the ACD+/- and ACD+/-WTA11 mutant uteri. Unsupervised clustering of the WT cells identified eight distinct cell types, including epithelial, endothelial, stromal, myocytes, lymphocytes, myeloid, pericytes and mesothelial. Subclustering was carried out for five of these cell types, for example dividing endothelial cells into capillary, arterial, venous and lymphatic cell subtypes. Epithelial cells were divided into glandular and luminal subtypes. In addition, we identified a considerable heterogeneity of stromal and myocyte cells. The results define the differential gene expression of these various cell types and subtypes and characterize the transcription factor codes that drive lineage specific development. The scRNA-seq data further allows the analysis of potential growth factor/receptor interactions between the different cell types in the PND12 uterus. These results confirm and significantly extend the previous laser capture microdissection/microarray study (Filant and Spencer, 2013) and scRNA-seq analysis of developing uterine epithelial cells (Wu et al., 2017).

As might be expected, the ACD+/- and ACD+/-WTA11 mutant uteri show the greatest gene expression changes in the myocyte and stromal compartments, where the mutated Hox genes are most strongly expressed. For these cell types the ACD+/-WTA11 cells show a striking intermediate gene expression state, between the ACD+/- and WT cells, consistent with their intermediate severity genotype (Fig. 9). There were also pronounced differences in gene expression of the ACD+/- mutant epithelial cells, even though they show very little expression of the mutated Hox genes, suggesting that this might be the result of disrupted cross talk with the mutant stroma.

Several growth factor/receptor pathways of interest were altered in the mutants. There was reduced *Gli1* and *Ptch2* expression in the ACD+/- uterus (Fig. 11), giving evidence for reduced hedgehog signaling. Both *Gli1* and *Ptch2* were also previously shown to be expressed in neonatal uterine stroma (Nakajima et al., 2012). There was also evidence of reduced Tgfbeta signaling in ACD+/- uteri. The mutant stromal cells show 2.5 FC reduced expression of thrombospondin (*Thbs1*), which functions to activate latent Tgfbeta (Sweetwyne and Murphy-Ullrich, 2012) and the epithelial cells show reduced expression (2.5 FC) of *Dcn*, which binds Tgfbeta and enhances its activity (Takeuchi et al., 1994).

Wnt signaling was dramatically reduced in the ACD+/- mutant uteri. Lef1 is a member the Tcf/Lef family of transcription factors and interacts with β-catenin to up-regulate Wnt target genes. Lef1 showed expression in a subset of stromal cells of WT PND10 uteri, especially in the mesometrial region (Fig. 11). Reduced Lef1 expression in ACD+/- mutants was shown by scRNA-seq, RT-PCR (Fig. 10) and IHC (Fig. 11). A majority of *Lef1*^*-/-*^ mice die by two weeks of age with missing teeth, mammary glands, whiskers, hair (Gendron et al., 1997) and uterine glands (Shelton et al., 2012). Taken together these results strongly suggest that the reduced expression of *Lef1* in ACD+/- mutants contributes to the failure of formation of normal numbers of uterine glands.

The ACD+/- mutant uteri stroma also showed reduced expression of *Ndp* (2.7 FC), which encodes Norrin, a non-Wnt ligand for Frizzled receptor, capable of activating canonical Wnt signaling (Xu et al., 2004). The mutants also showed lower expression of *Sp5* (Fig. 10) a known Wnt target (Huggins et al., 2017). These results provide additional evidence in support of diminished Wnt signaling in Hox mutants.

Previous studies have shown that Wnt signaling is essential for proper uterine gland formation. Wnt4 (Franco et al., 2011), Wnt7a (Miller and Sassoon, 1998), Wnt5a (Mericskay et al., 2004), Wnt11 (Hayashi et al., 2011) and beta catenin (Arango et al., 2005; Hernandez Gifford et al., 2009; Jeong et al., 2009) are all required for gland formation. The *Wnt7a* mutant is of particular interest. *Wnt7a* expression is in the luminal uterine epithelium postnatally, (Miller and Sassoon, 1998; Parr and McMahon, 1998) with mutants showing near complete absence of uterine glands, loss of postnatal expression of *Hoxa10* and *Hoxa11*, and partial homeotic transformation of uterus, surprisingly, towards vagina, as measured by the mutant stratified luminal epithelium, similar to the *Hox9,10,11* mutants we previously described (Raines et al., 2013). Further, *Hoxa11* is required to maintain *Wnt7a* expression (Mericskay et al., 2004). These combined observations suggest a positive feedback loop, with Hox genes activating Wnt signaling and Wnt signaling activating Hox genes. Of note, the ACD+/- uterus is not homozygous mutant for any Hox gene. The dramatic effect on Wnt signaling resulting from heterozygous mutation of nine related Abd-B Hox genes provides further evidence of their functional overlap.

The ACD+/- mutant uteri additionally showed greatly diminished (3.8 FC) stromal expression of *Cxcl12*, also known as stromal cell-derived factor 1 (*Sdf1*), which encodes a ligand for Cxcr4, expressed by the LE. In multiple developing contexts Cxcr4/Cxcl12 interaction has been previously shown to drive cell migration and tubulogenesis (Doitsidou et al., 2002; Feng et al., 2014; Ivins et al., 2015; Molyneaux et al., 2003; Ueland et al., 2009), key steps in uterine gland formation. Cxcl12 is a powerful chemoattractant for many cell types (Kim and Broxmeyer, 1999; Kulbe et al., 2004; Lazarini et al., 2003), and inward movement of LE cells is a key first step in uterine gland formation. Indeed, CXCL12 and its G protein-coupled receptor CXCR4 have been proposed to control cell movement in many developing organs (McGrath et al., 1999). For example, in the developing pancreas, similar to the uterus, *Cxcl12* is expressed in the mesenchyme while *Cxcr4* is expressed in epithelial cells. Either genetic mutation or pharmacologic inhibition of *Cxcl12* with AMD3100 resulted in a significant reduction of epithelial buds during pancreas branching morphogenesis (Hick et al., 2009). Similar, in the developing kidney the inhibition of Cxcl12/Cxcr4 with AMD3100 causes a severe reduction of ureteric bud branching morphogenesis (Ueland et al., 2009). Taken together these results give evidence for a role for the reduced mutant expression of *Cxcl12* in the observed formation of fewer uterine glands.

## Conclusions

In summary this study identifies a novel role for the *Hoxc9,10,11* genes in uterine gland formation. This function is redundant with the *Hoxa9,10,11* and *Hoxd9,10,11* genes and is only seen in a sensitized genotype with reduced expression of these paralogs. We further used scRNA-seq to define the gene expression patterns of the multiple cell types of the WT developing uterus. The results define the gene expression patterns driving lineage specific development. In addition, scRNA-seq was used to characterize the perturbed gene expression levels of all developing uterus cell types in the ACD+/- and ACD+/-WTA11 mutants. Particularly striking was the observed reduced Wnt signaling and disruption of the Cxcl12/Cxcr4 axis in the mutants.

## Materials and Methods

### Animals

All animal procedures were approved by the Institutional Animal Care and Use Committee at Cincinnati Children’s Hospital Medical Center according to their guidelines for the care and use of laboratory animals (IACUC protocol 2015-0065). The *Hox* mutant mice used in this study were generated by recombineering as previously described (Drake et al., 2018; Raines et al., 2013). The Hoxa9,10 mutant mice were made exactly as previously described for *Hoxa9,10,11*, using the same BAC targeting vector, but the recombination event inserted only the two *Hoxa9,10* mutant genes instead of all three.

The female reproductive tracts were isolated immediately after euthanasia using IACUCC approved methods, washed in cold 1X PBS, and processed for immunohistochemistry or the isolation of RNA using standard methodologies. For blastocyst isolation, gestational day 4 (vaginal plug = Day 1) uterine horns were gently flushed with 1 ml of 1X PBS and blastocysts counted using a dissecting microscope. For all experiments a minimum of 3 animals were analyzed per genotype.

### Hematoxylin and eosin (H&E) staining

Female reproductive tracts were isolated as described above, then fixed in 4% paraformaldehyde, rinsed in 1X PBS and dehydrated through ethanol washes. Uterine horns were cut into four pieces (ovary/oviduct and three uterine sections) and all pieces of tissue embedded into one paraffin block. Glands were counted in each transverse section from a minimum of three non-contiguous slides and the average number of glands per section determined.

### Immunohistochemistry

Briefly, fixed uterine horns were sectioned (5 μm), sections mounted on slides, baked, deparaffinized in xylene, and rehydrated in a graded series of ethanol. Antigen retrieval was conducted using 10 mM sodium citrate (pH 6.0) with boiling. Sections were blocked using 4% normal donkey serum in 1X PBST (pH 7.4) and incubated overnight at 4°C with a rabbit monoclonal anti-Lef1 (C12A5) antibody (Cell Signaling Technology) at dilution of 1:2000. Sections were washed in 1X PBST and incubated with biotinylated goat anti-rabbit secondary antibody (Vector Laboratories) at a dilution of 1:200 for 30 minutes at room temperature. After washes in 1X PBST, the immunoreactive protein was visualized using the Vectastain ABC kit (Vector Laboratories) and 3,3’ diaminobenzidine tetrahydrochloride hydrate (Sigma) as the chromagen. Sections were counterstained with nuclear fast red before coverslips were applied using Permount.

### RNA extraction and real-time PCR

RNA was isolated from uterine samples using the RNeasy Plus mini kit (Qiagen) after homogenizing the samples using Qiagen’s Tissue Lyser II. The quantity and purity of each RNA was determined using an Agilent Bioanalyzer nanochip. Total RNA (1 μg) was reverse transcribed to cDNA using the SuperScript Vilo kit (Invitrogen). The cDNA was ethanol precipitated and resuspended in nuclease-free water. cDNA concentration was determined using a NanoDrop Spectrophotometer and samples were stored at −20°C until use.

Real-time PCR was conducted using Applied Biosystems TaqMan Gene Expression Assays, TaqMan Gene Expression Master Mix, nuclease free water, and the Step One Plus Real Time PCR system (Thermo Fisher Scientific). *Gapdh* was used as the reference gene. All samples were analyzed in triplicate for each gene tested. Statistical analyses were performed using GraphPad Prism (Version 7).

### Cold protease uterine dissociation protocol for Drop-seq analysis

For *Bacillus licheniformis* enzyme mix 1 ml contained 890 ul of DPBS with no added Ca or Mg, 5 μl of 5mM CaCl_2_, 5 μl of DNase (Applichem A3778, 125 U), and 100 μl *Bacillus licheniformis* enzyme, 100mg/ml (P5380, Aldrich Sigma). Euthanized post-natal day 12 (PND12) mice (morning of birth = post natal day 0, PND0) and isolated uterine horns without ovary and oviduct. Incubated tissue in 1 ml of enzyme mix at 6°C for 2-4 minutes, then began to dissociate uterus by drawing tissue through an 18 gauge needle attached to a 3 ml syringe. Repeated 6°C incubation and passed tissue through progressively smaller gauge needles (23 and 21 gauge) until a single cell suspension was achieved. Progress was monitored with a microscope. When single cell status was achieved, added an equal volume of ice cold 10% FBS in 1X PBS to inactivate protease. Filtered cell suspension through a 30 μm MACS filter. Rinsed filter with 3-5 mls of ice cold 0.01% BSA in 1X DPBS into a 50 ml centrifuge tube. Transferred cells to a 15 ml centrifuge tube and pelleted cells by centrifuging at 160 g for 5 min at 4°C. Repeated washing and centrifugation steps twice more. Resuspended cells in 1-2 mls of 0.01% BSA in 1X DPBS and determine cell viability using Trypan blue. A hemocytometer was used to determine the final concentration of cells.

### scRNA-seq and data analysis

scRNA-seq for WT, ACD+/- and ACD+/-WTA11 was carried out using Drop-seq as previously described (Adam et al., 2017; Macosko et al., 2015).

Cell-type clustering and marker genes were identified using Seurat 2.3.2 (Butler et al., 2018) and R 3.4.3. Data were prefiltered at both the cell and gene level with the removal of cells with low library complexity (<500 expressed genes) as well as those with a high percentage (>20%) of unique molecular identifiers (UMIs) mapping to mitochondrial genes. In addition, cells with more than 8% histone gene expression and more than 0.4% hemoglobin gene expression were also excluded. Genes were selected that had > 0 UMI in more than five cells. Following filtering, the number of cells, the average number of genes called expressed per cell, and average number of transcripts (UMI) per cell respectively were, WT (6343, 1222, 2283), ACD+/- (2134, 1258, 2865), and ACD+/-WTA11 (3378, 1297, 2531). Genes with the highest expression variability were used for the principal component analysis. Significant principal components were determined using the PCElbowPlot function. Data were visually represented after clustering using Seurat’s t-SNE implementation. The FindAllMarkers function was used to determine the top marker genes for each cluster with a minimum of 10% of cells expressing the gene within the cluster and a minimum logFC threshold of 0.25. Sub clustering was performed by repeating the above analysis on individual clusters.

Differential expression between clusters was determined using the FindMarkers function in Seurat with pseudocount set to 0.01. Similar cell types from the WT experiment were compared to each of the two mutant mice.

The top differentially expressed genes were filtered on genes that had a log2 fold-change greater than 1 and were expressed in at least 10% of the cells for that cluster.

Data has been deposited in GEO under accession number GSE118180.

#### Circos plot

The Circos plot was generated by identifying the growth factors and receptors that are expressed in greater than 7.5% of the cells within any given cluster. The possible interactions between the stromal growth factors and their corresponding receptors in the multiple cell types were plotted. The Circos plot was generated as previously described (Krzywinski et al., 2009).

#### Growth factor and receptor analysis

The growth factors in the stromal groups and their corresponding receptors in the epithelial compartments were input into GeneMania to generate a network plot as previously described (Warde-Farley et al., 2010).

## Acknowledgements

This work was supported by NIH grant RO1 DK099995 to SSP.

